# The Mechanisms of GPR55 Receptor Functonal Selectivity

**DOI:** 10.1101/2022.01.12.476071

**Authors:** Mikhail G. Akimov, Natalia M. Gretskaya, Polina V. Dudina, Galina D. Sherstyanykh, Galina N. Zinchenko, Oksana V. Serova, Ksenia O. Degtyaryova, Igor E. Deyev, Vladimir V. Bezuglov

## Abstract

The objective of the study was to establish the mechanisms of multidirectional signal transmission through the same G-protein coupled receptor GPR55. Using the CRISPR-Cas9 system, clones of the MDA-MB-231 line knockout for the GPR55, CB1, CB2, GPR18, and TRPV1 receptor genes were obtained. On clones of the MDA-MB-231 line with a knockout CB2 receptor, the cytotoxic activity of the pro-apoptotic ligand docosahexaenoyldopamine (DHA-DA) did not change or slightly increased, while the pro-proliferative activity of the most active synthetic ligand of the GPR55 receptor (ML-184) completely disappeared. On the original line MDA-MB-231, the stimulatory effect of ML-184 is removed by the CB2 receptor blocker, but not by GPR55. At the same time, the stimulating effect of ML-184 is practically not manifested on cell lines knockout at the GPR55 receptor. Thus, it can be confidently assumed that when proliferation is stimulated with the participation of the GPR55 receptor, a signal is transmitted from the CB2 receptor to the GPR55 receptor due to the formation of a heterodimer. GPR18 and TRPV1 receptors are additionally involved in the implementation of the cytotoxic effect of DHA-DA, while the CB1 receptor is not involved. In the implementation of the cytotoxic action of DHA-DA, the predominant participation of one of the Gα subunits was not found, but the Gα_13_ subunit plays a decisive role in the implementation of the proproliferative action.

## 1. Introduction

GPR55 is a G protein-coupled receptor. Initially, it was assumed that its ligands were endocannabinoids (anandamide, 2-arachidonoylglycerol, etc.) and is still considered a non-classical cannabinoid receptor. Later, it was found that the active stimulator of this receptor is lysophosphatidylinositol (LPI), which is considered the primary natural ligand [1] of the GPR55 receptor. We have shown that another natural ligand for this receptor can be arachidonoyldopamine, as well as other acyl dopamines [2,3]. Thus, the GPR55 receptor is currently positioned as a multi-ligand receptor.

Evidence from pharmacological studies of the GPR55 receptor indicates that its signaling and corresponding cellular responses depend on the type of ligand that activates it. For such receptors, the concept of ‘agonist functional selectivity’ has been proposed [4].

The GPR55 receptor is expressed in many organs and tissues of the body [5]. In particular, transcripts of this receptor have been found in the central nervous system (the striatum, hypothalamus, microglia, etc.). This receptor is present in the spleen, bone marrow, platelets, neutrophils, and lymphocytes, which indicates its involvement in the inflammatory response. The GPR55 receptor has been identified in various types of cancer cells (glioma, melanoma, breast and pancreatic cancer [6].

Activation of GPR55 LPI leads to increased cell proliferation, and therefore, if an increased level of expression of this protein is detected in tumors, this is considered a marker of a poor prognosis for the development of the disease [7].

On the other hand, activation of GPR55 by anandamide and acyl dopamines induces cell death via apoptosis. The exact mechanism of implementation of this process may differ: for example, in the cells of rat pheochromocytoma PC12, the expression of NO synthase is stimulated, which synthesizes reactive oxygen species as a by-product [3]. In the case of anandamide in cholangiocarcinoma cells, the death receptor is activated with the participation of GPR55 [8].

Literature data on GPR55 intracellular signaling indicate that this receptor, depending on the cell line and conditions, is able to interact with several G alpha subunits: Ga13, Gaq/11, Ga12, Ga12/13 (Morales, Reggio, Cann. Cann. Res., 2017), while the Gai1/2, Gai3, and Gas subunits do not interact with this receptor (Ross and TIPS, 2009).

The potential for interaction with several Ga subunits has also been described for other G protein-coupled receptors, for example, after activation of the LPA4 receptor, neurite retraction mediated by Ga12/13 and Rho, calcium mobilization with the participation of Gaq, and an increase in cAMP levels with the participation of Gas (Ross, TIPS, 2009). At the same time, in the case of this receptor, the activation of different G alphas can be due either to different isoforms of the receptor, or to the fact that only a part of the G alpha subunits is present in the cells.

Two of the mentioned subunits lead to the release of intracellular calcium: activation of Gq leads to the inclusion of phospholipase C and the accumulation of IP3, a calcium channel ligand on the ER IP3R, and Ga12 stimulates an additional Rho protein, which can also stimulate phospholipase C and, in addition, the actin cytoskeleton. Calcium release is an intermediate event in the activation of ERK and NFAT by the GPR55 receptor (Lauckner et al., PNAS, 2008).

The interaction of GPR55 with different pathways within the cell varies depending on the ligand. Thus, in HEK 293 cells transfected with GPR55, LPI, the best known agonist of this receptor, causes intracellular calcium release, NFAT activation, ERK1/2 phosphorylation, CREB and NFkB activation. At the same time, treatment with a CB1 receptor antagonist (without LPI) has a similar effect on calcium and CREB, even more effective in the latter case, but suppresses ERK phosphorylation and NFkB activation (Henstridge et al., Br.J. Pharmacol, 2010).

The activity of GPR55 ligands depends on the concentration in a non-standard way: there is evidence that low concentrations of anandamide activate the receptor, while high concentrations inhibit it (Sharir et al., J.Neuroimmun.Pharmacol, 2012).

The formation of dimers between GPR55 and cannabinoid receptors CB1 and CB2 has been described. At the same time, the presence of CB1 reduces ERK activation caused by GPR55 ligands, while the presence of GPR55, on the contrary, enhances the activation of this kinase by CB1 ligands (Kargk et al., JBC, 2012). Upon dimerization of GPR55 with CB2, the first receptor suppresses the activity of the second one (Moreno et al., JBC, 2014). In addition to the above-mentioned stimulation of cancer cell proliferation and induction of apoptosis, GPR55 is also involved in other processes. For example, activation of this receptor protects neurons from excitotoxicity in a model of cultured hippocampal slices (Kallendrusch et al., Glia, 2013). Alpha-lysophosphatidylinositol, through the activation of GPR55, has a microglia-mediated neuroprotective effect (Kallendrusch et al., Glia, 2013) and induces differentiation of retinal cells (Cherif., et al., ENeuro, 2015).

Most of the available literature describes GPR55 as a receptor associated with poor tumor prognosis, since its activation by lysophosphatidylinositol leads to increased proliferation of transformed cells. At the same time, there are a number of articles, both by the authors of the project and by foreign researchers, indicating that activation of the same receptor by acyl dopamine and anandamide induces apoptosis of tumor cells. Currently, there is no understanding of what mechanism provides for the transmission of such different signals by different GPR55 ligands through the same receptor.

The aim of the project is to investigate what mechanisms provide the ability of the non-classical cannabinoid receptor GPR55, depending on the structure of the ligand, to induce both proliferation and apoptosis of cancer cells when activated by endogenous lipids. Based on the known features of signaling through the GPR55 receptor, we suggest the possibility of two main mechanisms of the effect of the multidirectional action of the ligands of this receptor:

1. Ligands can induce dimerization or bind to various dimers of the GPR55 protein and CB1 or CB2 receptors (GPR55-CB1; GPR55-CB2) and thereby induce various intracellular signaling cascades.
2. Ligands of types A or P can bind to the GPR55 receptor coupled to different G-alpha subunits (q, 12 or 13), or provide preferential interaction of the receptor with a certain set of G-alpha subunits, which leads to opposite consequences for the cell (apoptosis or proliferation).

## 2. Results

### 2.1. GPR55, GPR18, CB1, CB2, and TRPV1 knockouts generation

#### 2.1.1. GPR55

Restriction of the PCR fragment showed the absence of the NcoI restriction site in the modified DNA fragment. Individual cell clones were then obtained by multiple dilution and genomic DNA was isolated from the obtained clones. The PCR fragment containing the modified gene region was cloned into the pAL2-T vector; sequencing confirmed the presence of deletions in the modified gene in different alleles in C12 and F10 cell clones.

On the example (figure) there was a deletion of two nucleotides, leading to a premature stop codon. Thus, the MDA-MB-231 cell line knockout for the GPR55 gene was obtained.

**Figure 1.**
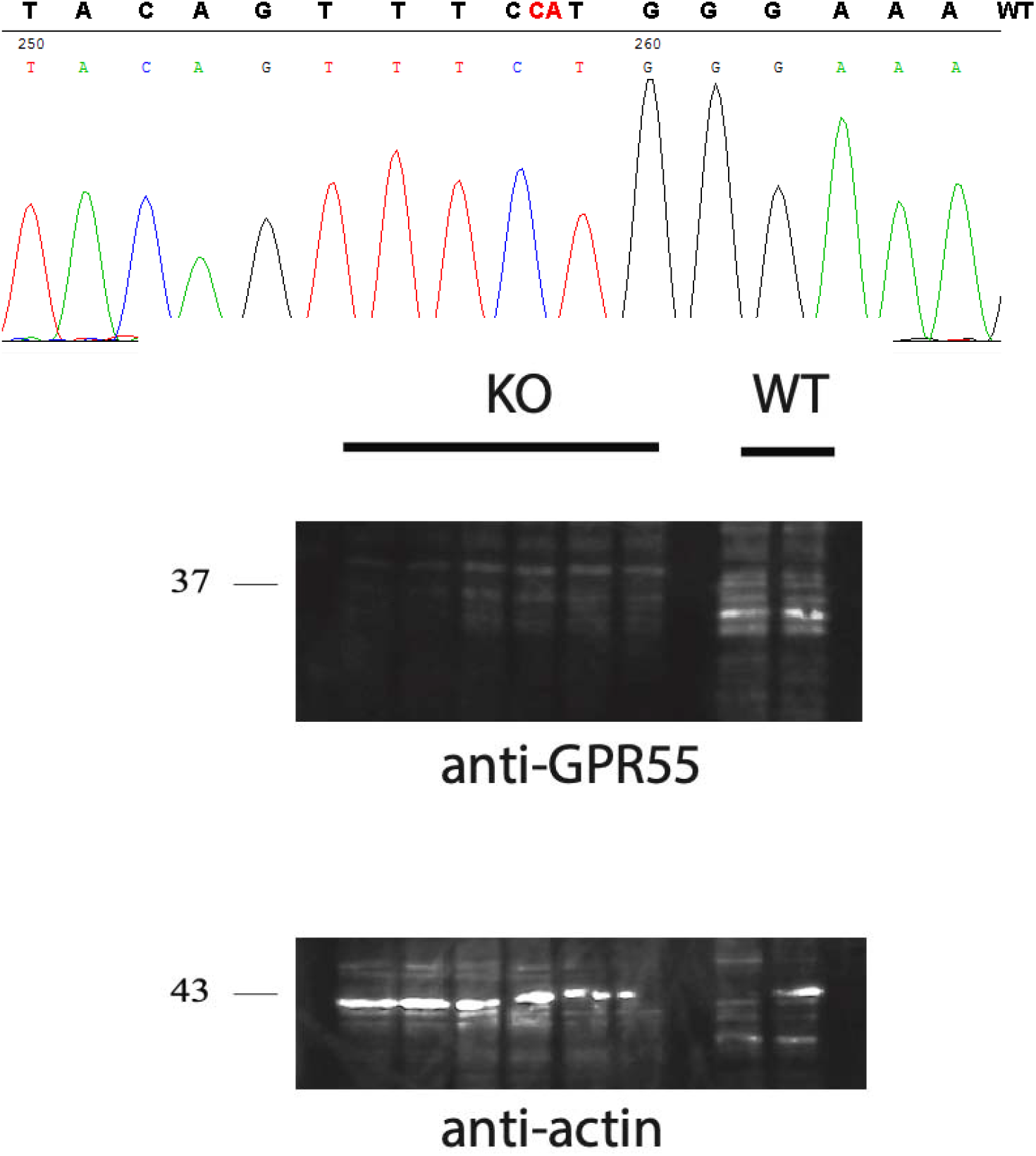
Sequence of a PCR fragment containing a modified region of the GPR55 gene. Above is a fragment of the wild-type GPR55 sequence. Below are Western blot data.

#### 2.1.2. CNR1 and CNR2

CNR1 genomic DNA contained a PstI restriction site at the Cas9 cleavage site, and PasI in the case of CNR2. Sequencing of the obtained plasmids confirmed the presence of the target insert. Restriction of the PCR fragment showed the absence of the PvuII restriction site in the modified DNA fragment in one of the three cell clones, indicating that the target gene had been modified in these cells. The genomic DNA fragment in cell clones 4 and 5 was partially cleaved with PvuII restriction enzyme.

In the case of CNR1, the PstI restriction site was preserved, indicating that the gene modification did not occur. Staining of cell lysates with antibodies specific for CNR1 using Western blot showed the presence of the protein in these cells. Obtaining individual clones was not carried out.

Analysis of the CNR2 gene fragment revealed the absence of the PasI restriction site, which indicated that the target gene had been modified. Individual cell clones were then obtained by multiple dilution and genomic DNA was isolated from the obtained clones. The PCR fragment containing the modified gene region is cloned into the pAL2-T vector. Sequencing confirmed the presence of deletions in the modified gene in different alleles in cell clones 2F1, 1H6, 1H9, 1B3, 2E5, 1B10. In the example, clone 2F1 had a single nucleotide deletion resulting in a premature stop codon. Thus, the MDA-MB-231 cell line, knockout for the CNR2 gene, was obtained.

**Figure 2.**
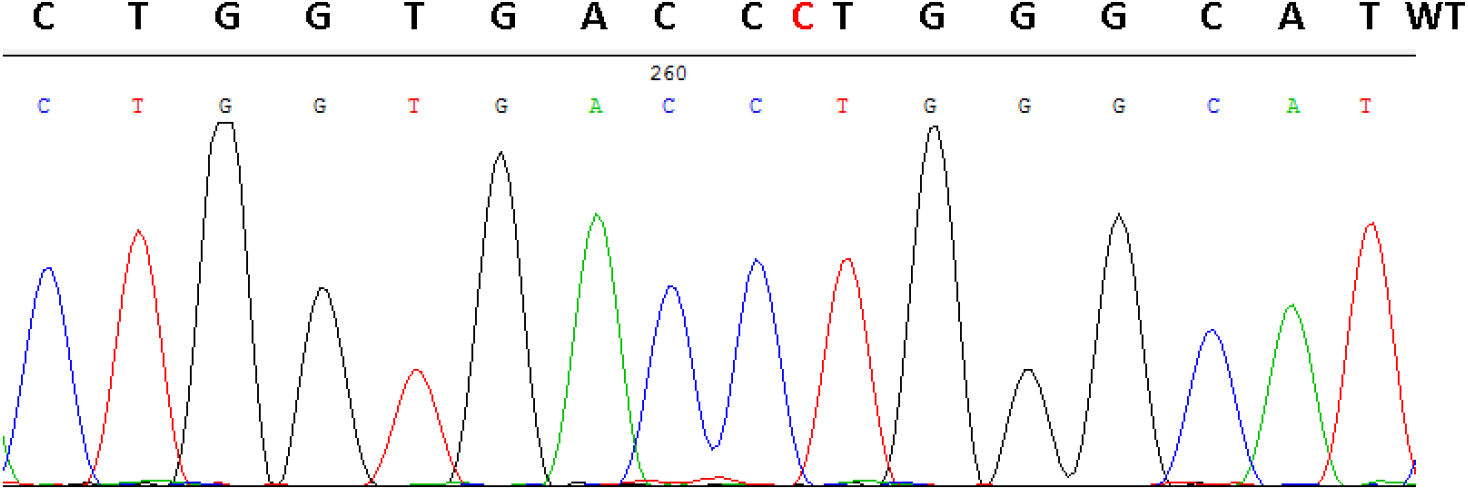
Sequence of a PCR fragment containing a modified region of the CNR2 gene, clone 2F1. Above is a fragment of the wild-type CNR2 sequence.

**Figure 3.**
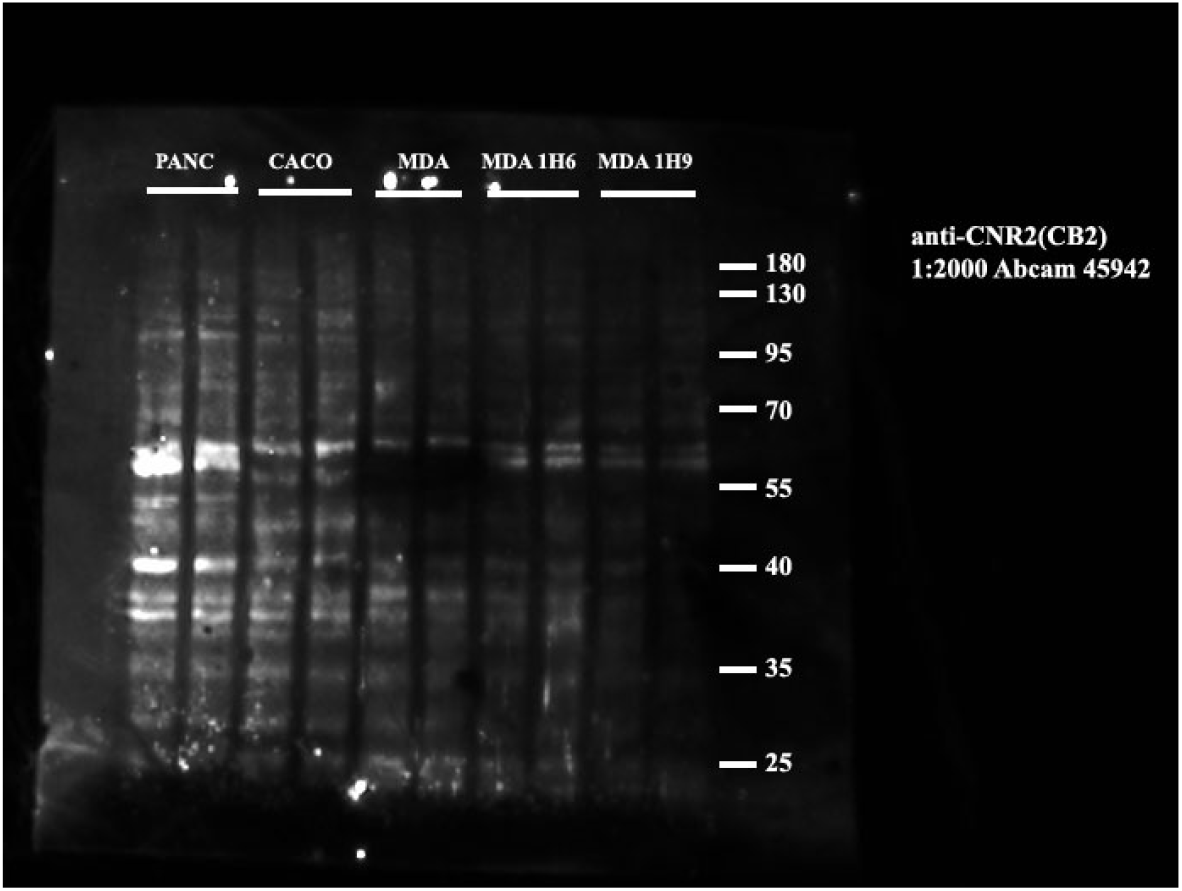
Detection of CB2 receptor expression in different cell lines, including two knockout lines, by Western blot (target molecular weight 40 kDa)

**Figure 4.**
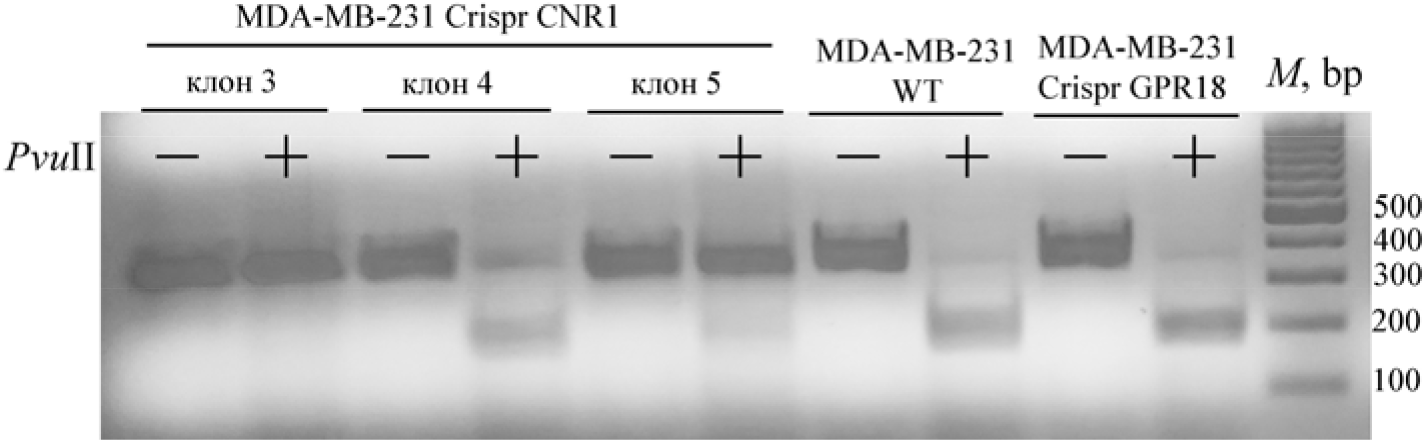
Electrophoresis of CNR1 genomic DNA fragments obtained by PCR before (-) and after (+) PvuII restriction enzyme treatment.

#### 2.1.3. TRPV1

At this stage of the work, we encountered the problem that the TRPV1 genomic DNA fragment of the original MDA-MB-231 cell line was not completely cleaved with the NcoI restriction enzyme, which indicates a possible replacement of nucleotides in this part of the genomic DNA. The genome region we have chosen is only partially suitable for obtaining a knockout cell line for the TRPV1 gene.

**Figure 5.**
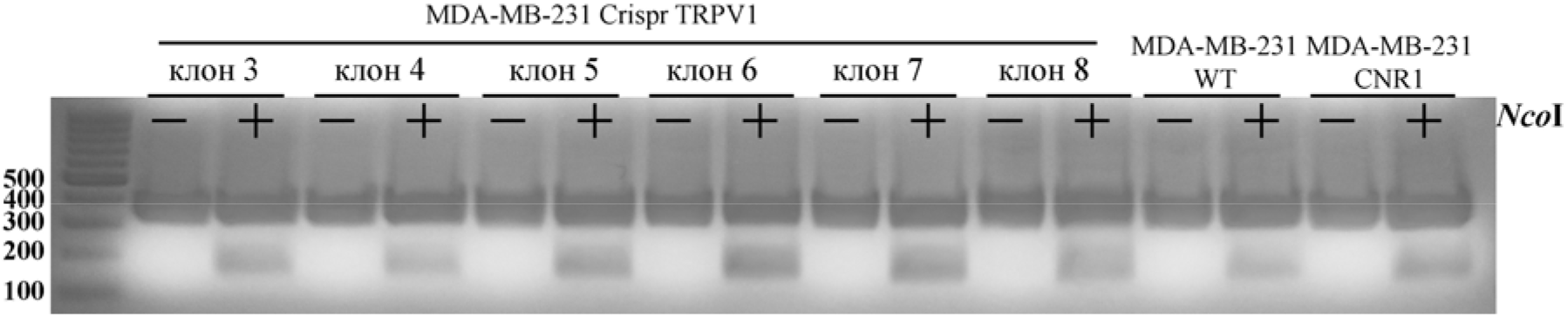
Electrophoresis of TRPV1 genomic DNA fragments obtained by PCR before (-) and after (+) treatment with NcoI restriction enzyme.

### 2.2. CB1 Knockout Effect

**Figure 6.**
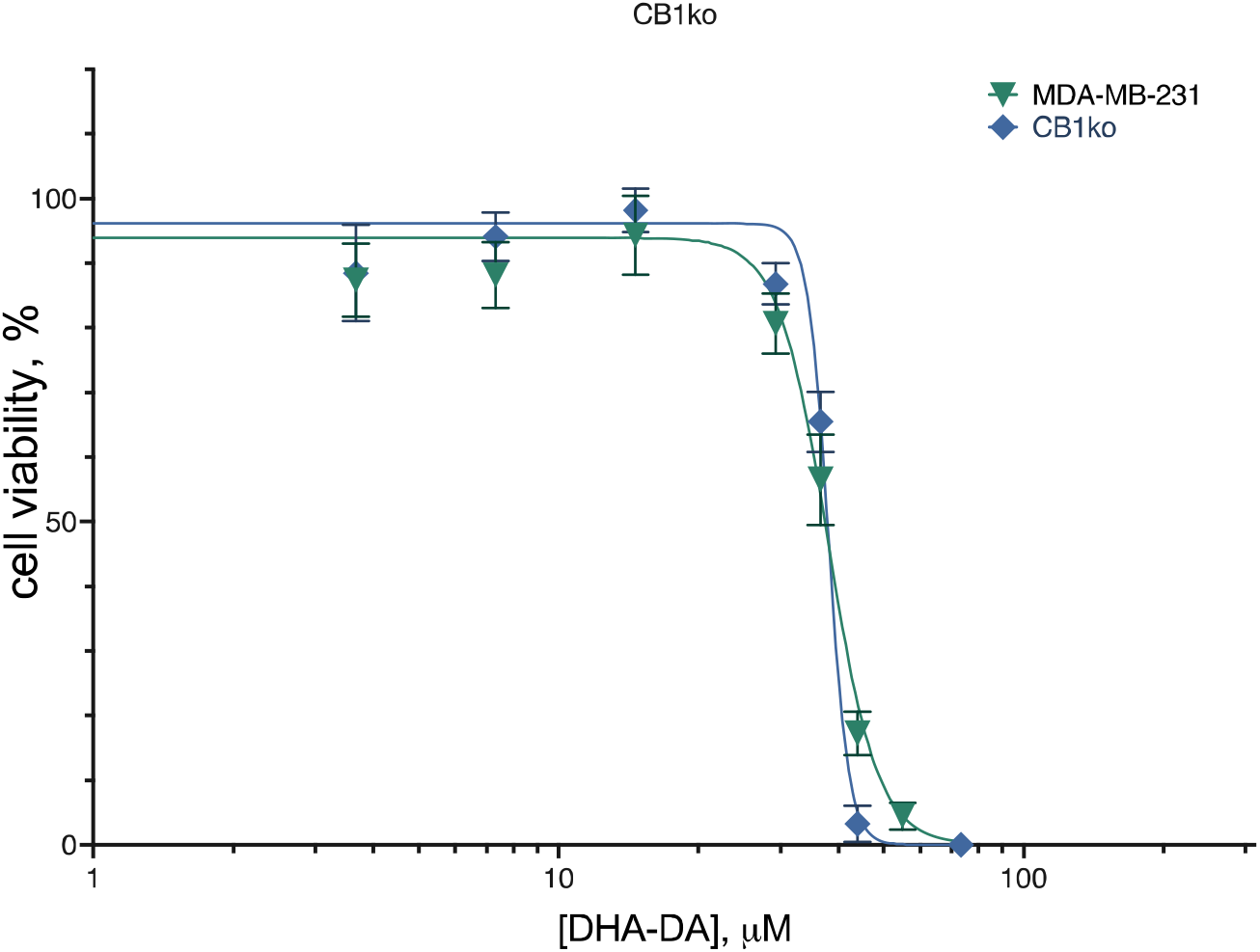
Effect of CB1 knockout on DHA-DA cytotoxicity. Incubation 24 hours, MTT test, mean±standard deviation (N=7 experiments)

**Table 1.**
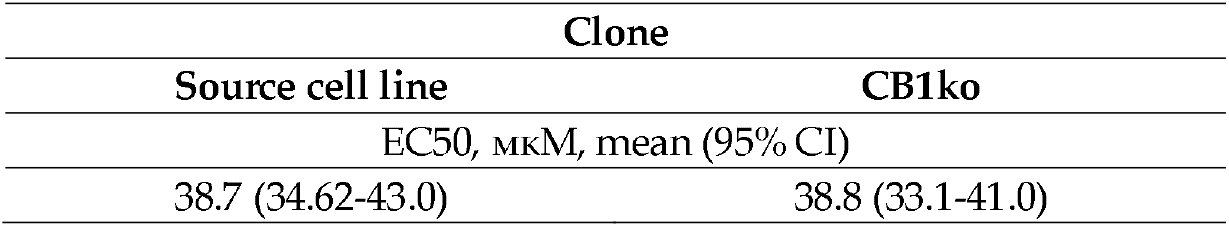

Knockout of the CB1 receptor did not lead to a significant change in the cytotoxicity of DHA-DA, that is, this receptor either does not participate in the implementation of the cytotoxic effect of the substance, or its contribution is insignificant and is compensated by the activity of other cannabinoid receptors.

### 2.2. CB2 Knockout Effect

The work was devoted to testing the role of the GPR55-CB2 heterodimer. We used 6 clones of the MDA-MB-231 cell line, knockout for the CNR2 gene, and 3 clones of the same line, knockout for the GPR55 gene. To improve reproducibility, seeding and treatment with the substance for all clones were carried out simultaneously.

DHA-DA was cytotoxic to both the original cell line and all clones with a knockout CB2 receptor. At the same time, for the original cell line, as well as for clones 1h9 and 2e5, the EC50 values were in the range of 37-41 µM, and for the remaining clones, in the range of 32–34 µM, and this difference was statistically significant. According to the sequence analysis of clones, it was in those clones where an increase in DHA-DA cytotoxicity was observed that the expression of the CB2 receptor was disrupted.

Thus, the removal of the CB2 receptor did not lead to a decrease in toxicity, and, therefore, the CB2-GPR55 dimer is not involved in this type of activity. However, the results of an increase in DHA-DA cytotoxicity after removal of the CB2 receptor provide indirect evidence for the interaction of these two G-protein coupled receptors.

**Table 2.**

| Clone |  |  |  |  |  |  |
| --- | --- | --- | --- | --- | --- | --- |
| MDA-MB-231 | 2f1 | 1h6 | 1h9 | 1b10 | 1b3 | 2e5 |
| EC50, μM, mean (95% C.I.) | | | | | | |
| 40.58 | 32.98 | 32.84 | 38.88 | 30.38 | 33.57 | 37.28 |
| (38.13 to 43.12) | (30.40 to 35.69) | (28.27 to 37.84) | (36.34 to 41.53) | (26.95 to 34.06) | (31.35 to 35.88) | (34.69 to 39.99) |

Comparison of the activity of ML184 on the original cell line and on lines knocked out at the CB2 receptor revealed that stimulation of proliferation was no longer observed on knockouts, which suggested the likely involvement of the CB2 receptor in the stimulation of proliferation with the participation of GPR55 agonists.

**Figure 7.**
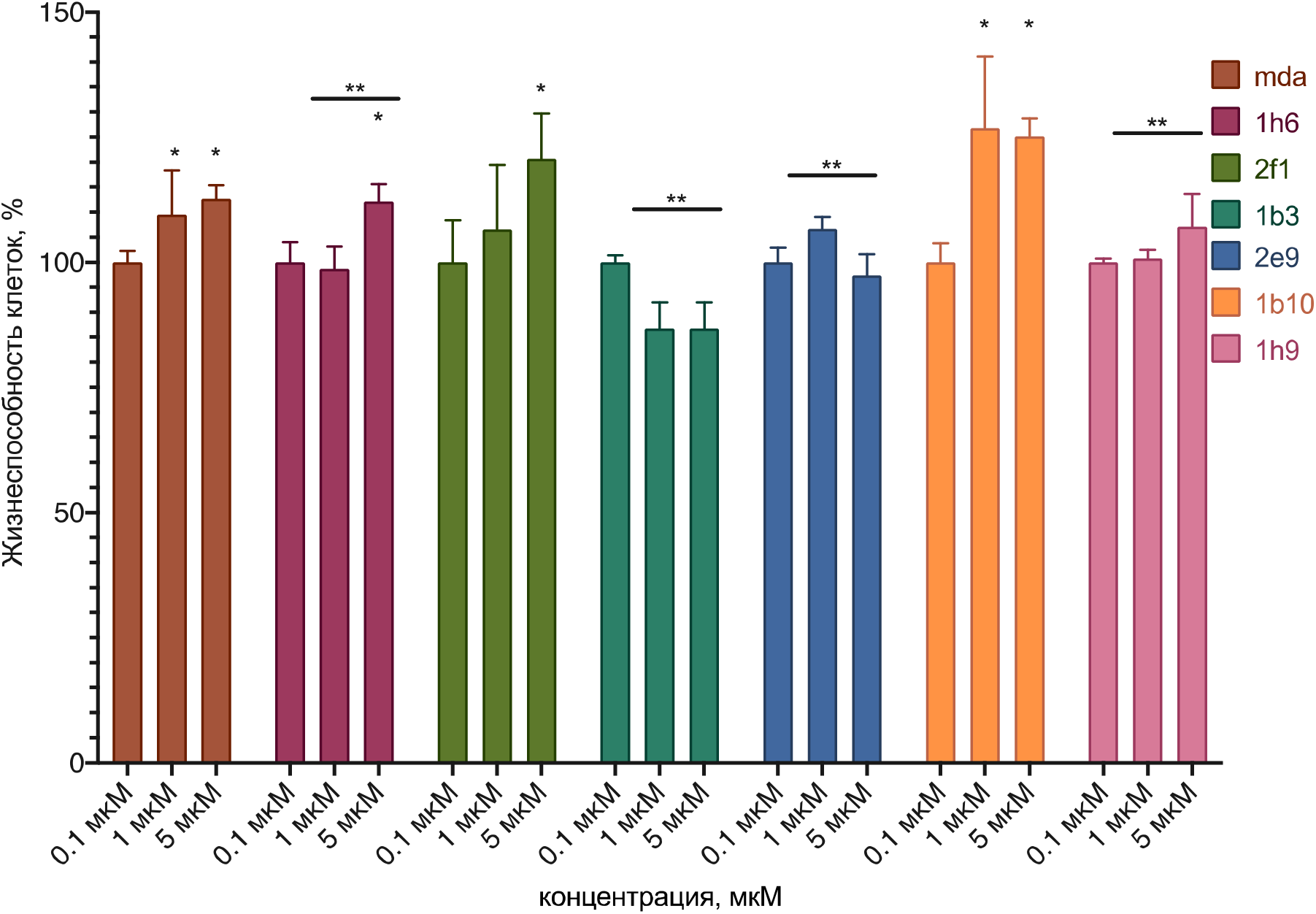
Comparison of ML184 activity on the original cell line and on lines knocked out at the CB2 receptor. *, statistically significant difference from control without substance; **, statistically significant difference from control without blocker

The fixed stimulation of proliferation and its disappearance could be interpreted in several ways: side interaction of the ligand with the CB2 receptor without the participation of GPR55; signal transmission in the GPR55-CB2 complex; damage in the process of knockout of other cell systems that are not related to the direct interaction of CB2 and GPR55.

To clarify the true state of affairs, the action of the ML-184 agonist was tested on the original cell line against the background of GPR55 (ML-193) and CB2 (SR 144528) blockers, as well as on the MDA-MB-231 cell lines knockout for the GPR55 receptor. It was found that the stimulatory effect of ML-184 on the original cell line was removed by the CB2 receptor blocker, but not by GPR55. At the same time, the stimulating effect of ML-184 is practically not manifested on cell lines knockout at the GPR55 receptor.

**Figure 8.**
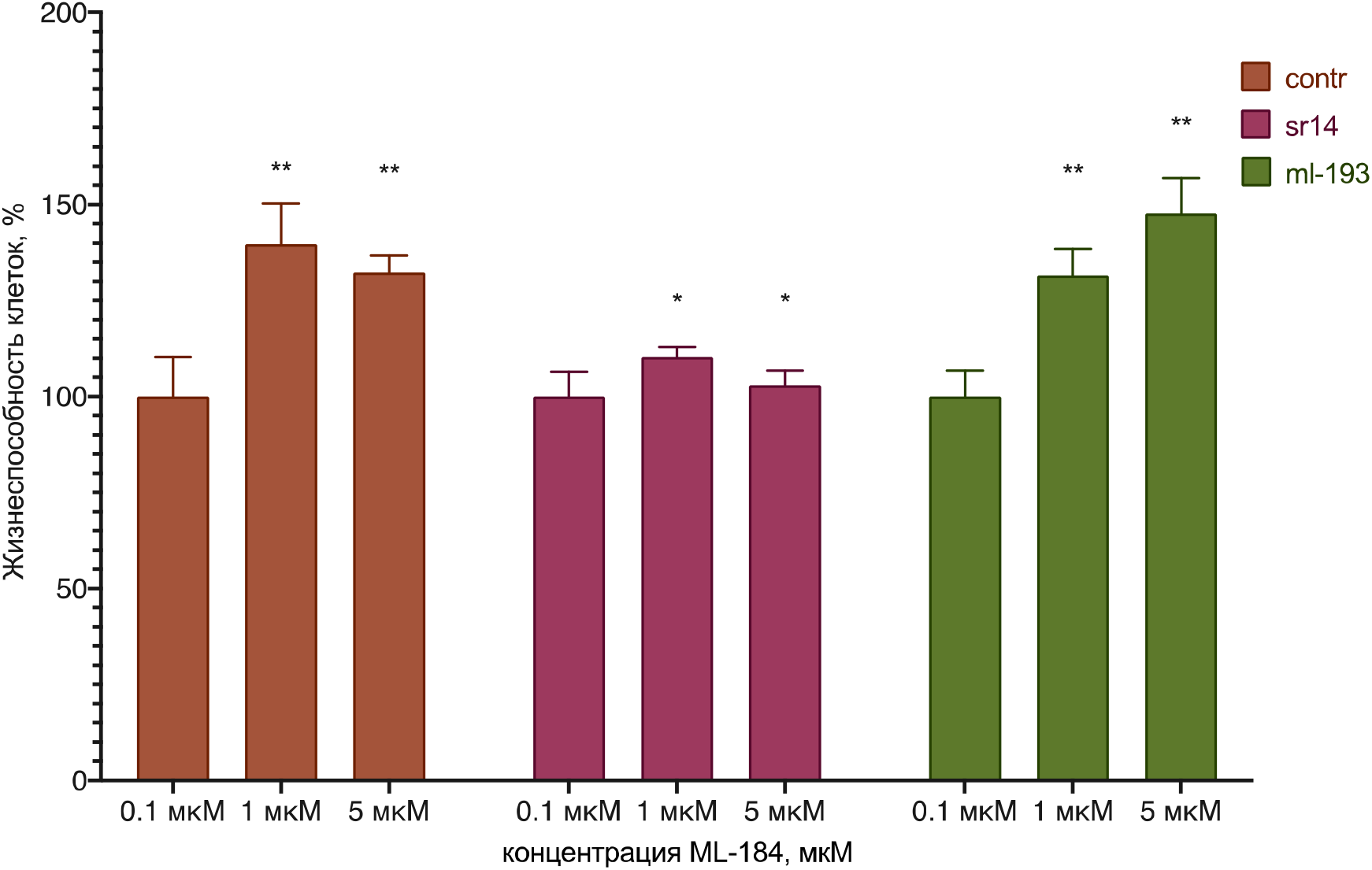
The effect of ML-184 on the original cell line against the background of GPR55 (ML-193) and CB2 (SR 144528) blockers. *, statistically significant difference from control without substance; **, statistically significant difference from control without blocker

**Figure 9.**
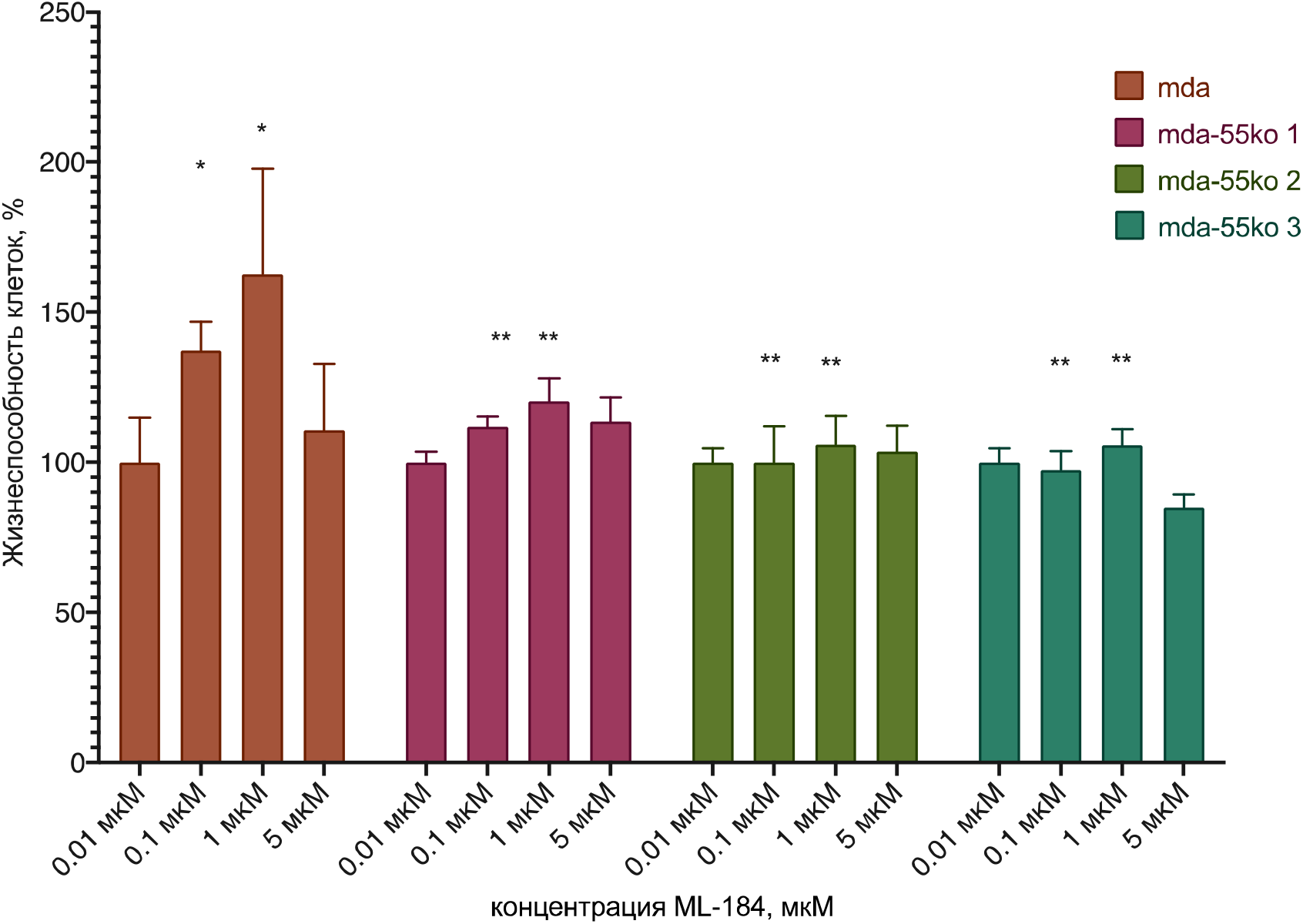
Effect of ML-184 on MDA-MB-231 cell lines knockout for the GPR55 receptor. *, statistically significant difference from control without substance; **, statistically significant difference from the original cell line

Thus, it can be safely assumed that stimulation of proliferation with the participation of the GPR55 receptor results in signal transduction from the CB2 receptor to the GPR55 receptor with the formation of a heterodimer.

### 2.3. TRPV1 Knockout Effect

**Figure 10.**
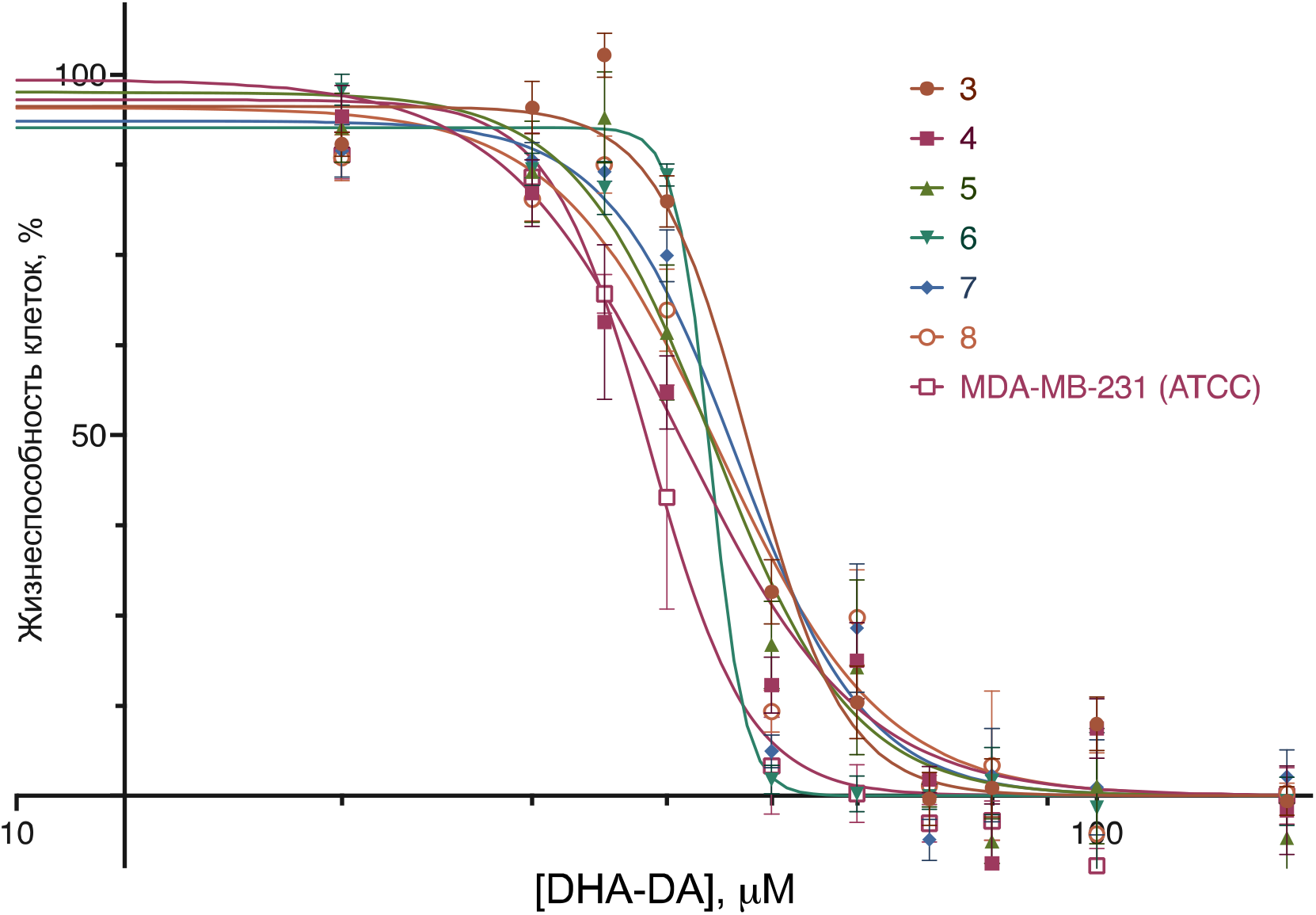
Cytotoxicity of DHA-DA for the original line MDA-MB-231 and clones with knockout TRPV1, incubation time 24 hours, MTT test data. Pooled data from 8 experiments, mean±standard error

**Table 3.**

| MDA-MB-231 | Knockout Clone |  |  |  |  |  |
| --- | --- | --- | --- | --- | --- | --- |
|  | 3 | 4 | 5 | 6 | 7 | 8 |
| EC50 (95% CI), uM |  |  |  |  |  |  |
| 38.55<br>(36.54-40.54) | 47.83*<br>(45.48-50.29) | 41.22<br>(38.58-44.09) | 44.14*<br>(41.11-47.52) | 43.89*<br>(41.21-47.72) | 46.47*<br>(43.97-49.27) | 44.75*<br>(42.08-47.58) |

\*, statistically significant difference from the original cell line, p<0.05, ANOVA with Holm-Sidak post-test

All clones recorded an EC50 value higher than that of the original cell line, i.e. DHA-DA toxicity decreased. The decrease was small (no more than 5-10 µM), but statistically significant; it was most pronounced in clones 3 and 7.

Taking into account the sequencing data and literature data, it can be assumed that the toxicity of DHA-DA in the considered cell line is partially realized through the TRPV1 receptor; however, the contribution of this receptor and the presence of its interaction with GPR55 require further study.

### 2.4. GPR18 Knockout Effect

**Figure 11.**
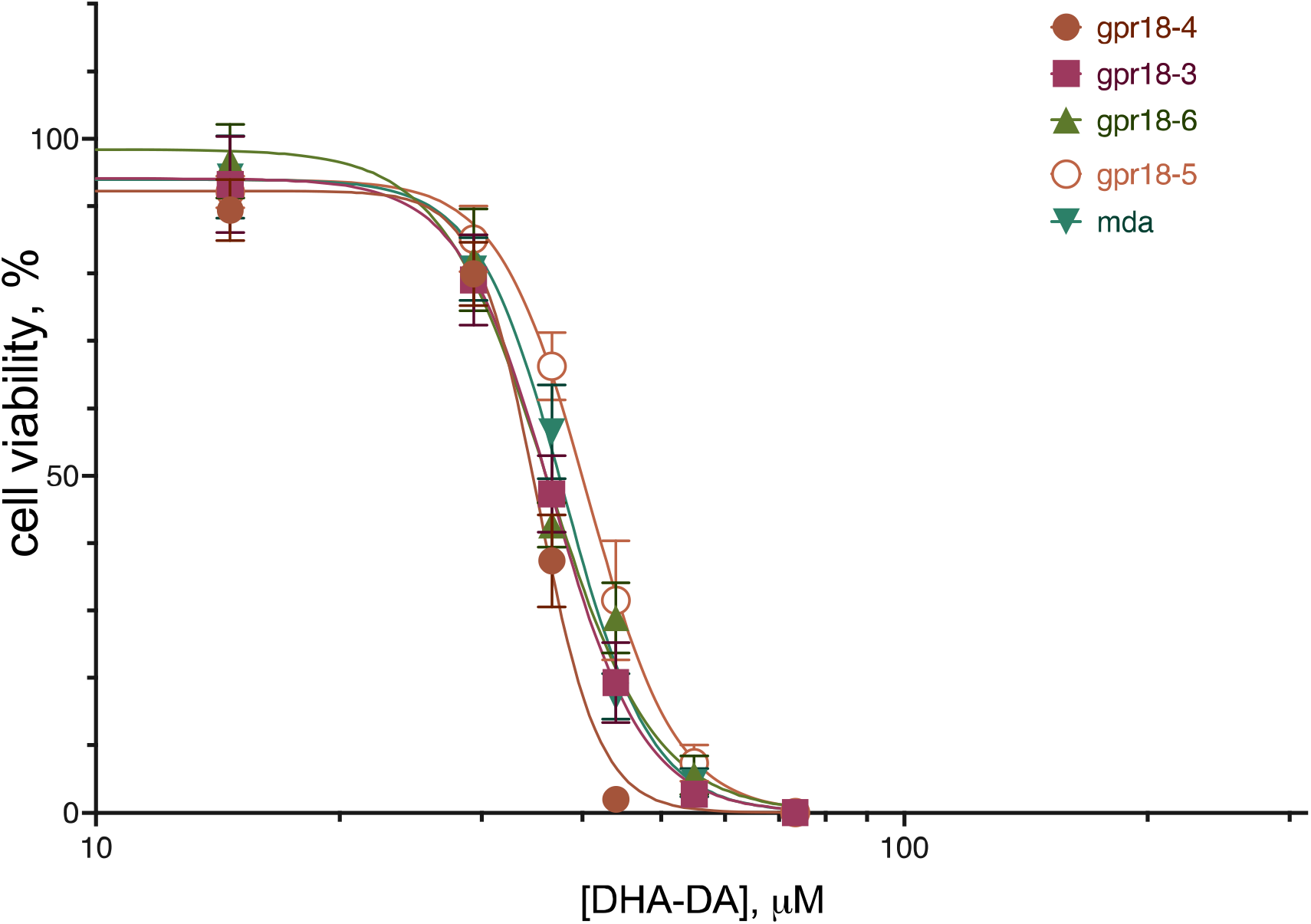
Cytotoxicity of DHA-DA for the original line MDA-MB-231 and variants of lines with knockout GPR18, incubation time 24 hours, MTT test data. Pooled data from 7 experiments, mean±standard error

**Table 4.**
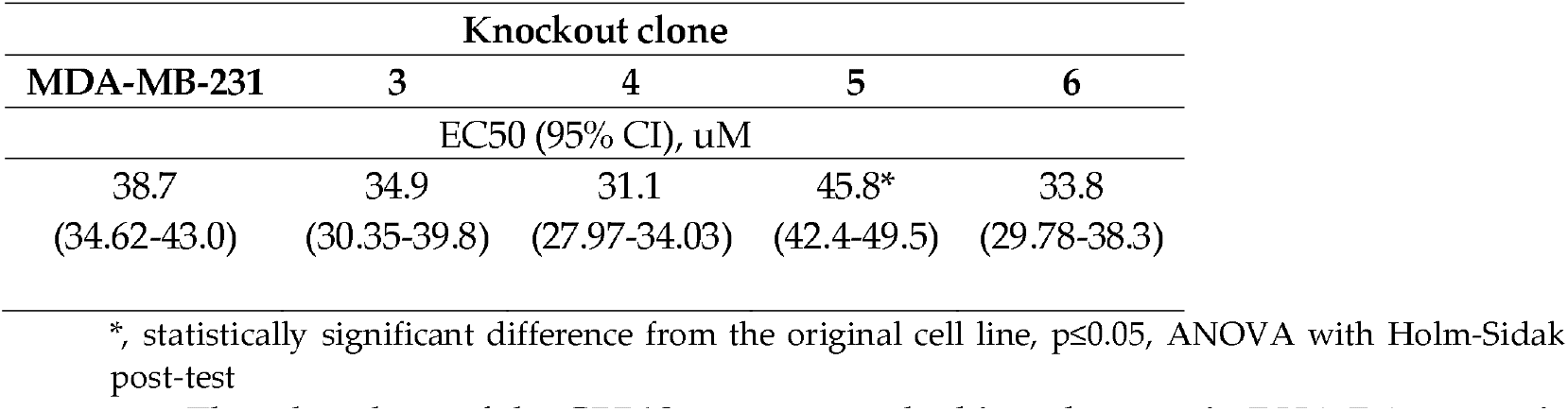

Thus, knockout of the GPR18 receptor resulted in a decrease in DHA-DA cytotoxicity for one of the obtained knockout populations and for all of its individual clones. This indicates the participation of this receptor in the implementation of the cytotoxic effect of the substance, however, the nature of this participation (parallel signal transduction or dimerization) requires further study.

### 2.5. G Protein Knockdown Effect

**Figure 12.**
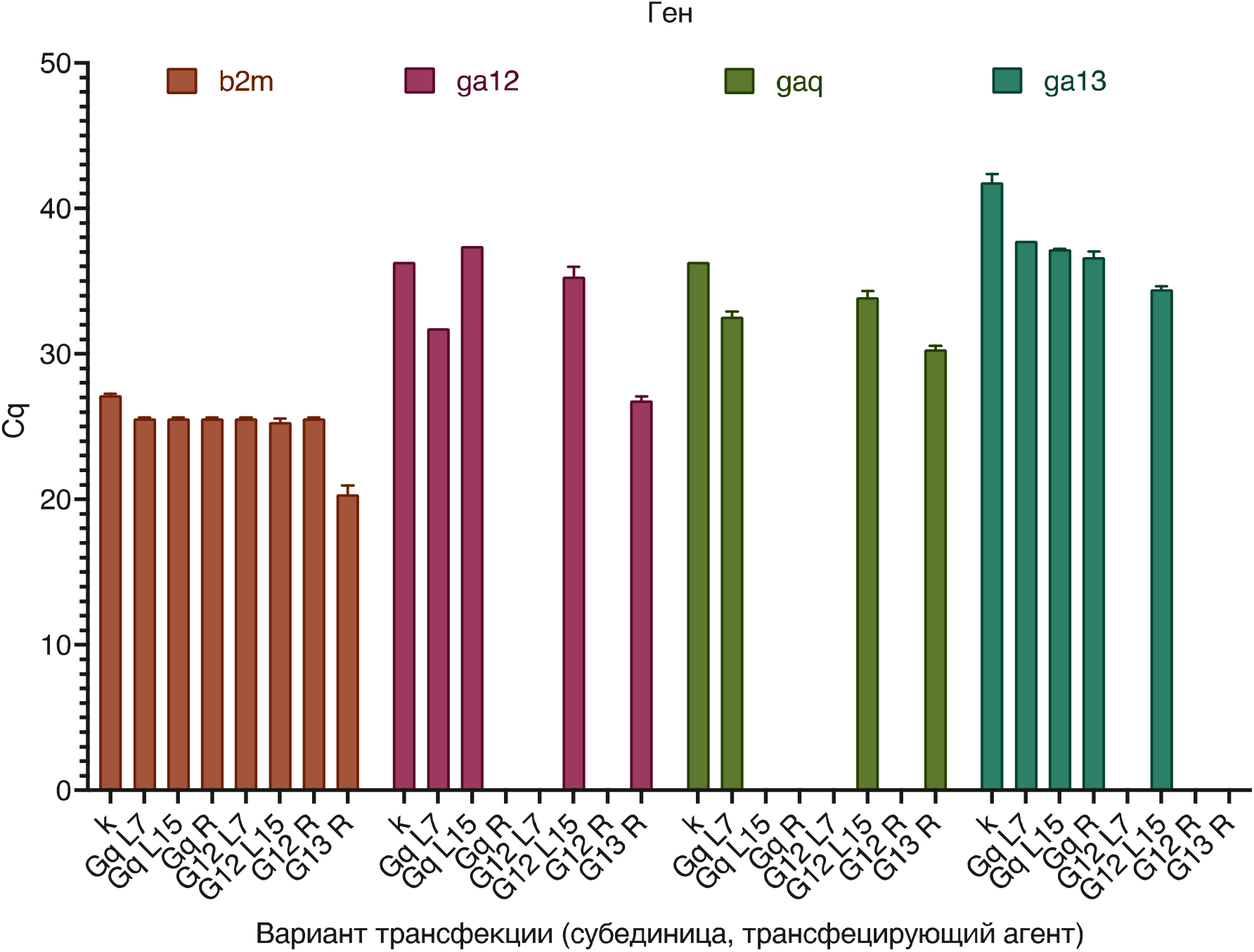
Expression of mRNA of subunits Gα_12_, Gα_13_ and Gα_q_ in MDA-MB-231 cells before and after siRNA knockdown with different variants of the transfecting agent (along the X axis L7 – lipofectamine 0.75 µl, L15 – lipofectamine 1.5 µl, R – RNAiMax). RT-qPCR data, Cq±SEM (higher value=lower expression)

**Figure 13.**
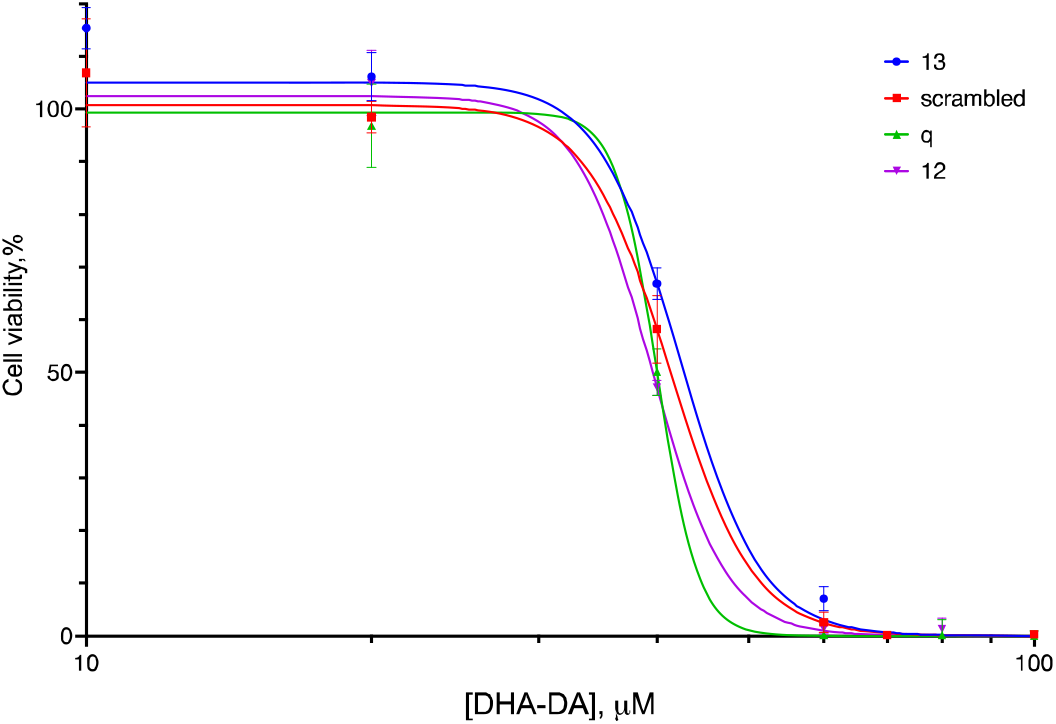
Cytotoxicity of DHA-DA for the MDA-MB-231 line against the background of knockdown of Gα subunits. Neq - scrambled control. MTT test data, mean±standard deviation

**Table 5.**
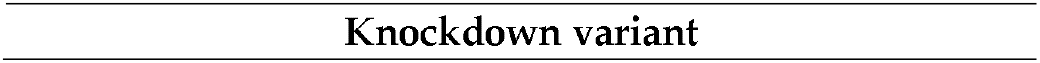
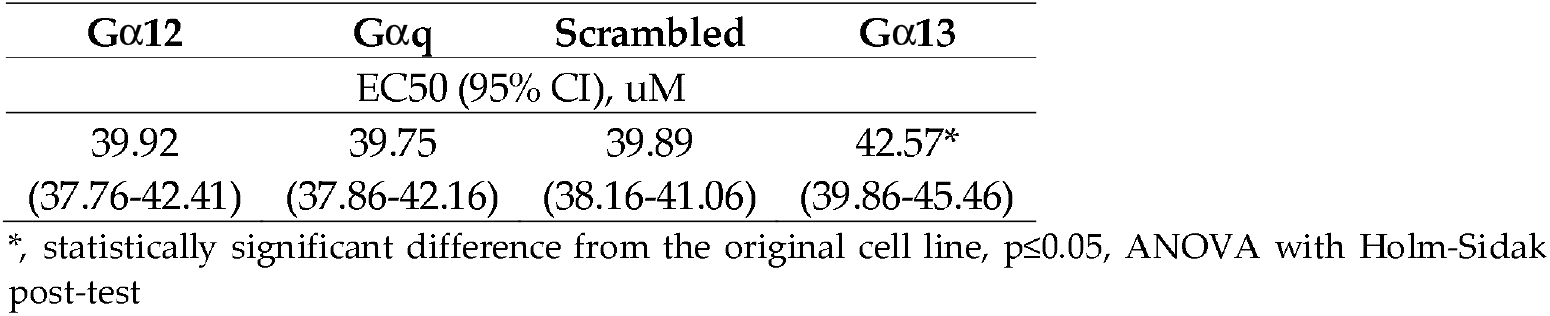

| Knockdown variant |
| --- |

| Gα12 | Gαq | Scrambled | Gα13 |
| --- | --- | --- | --- |
| EC50 (95% CI), uM |  |  |  |
| 39.92<br>(37.76-42.41) | 39.75<br>(37.86-42.16) | 39.89<br>(38.16-41.06) | 42.57*<br>(39.86-45.46) |

\*, statistically significant difference from the original cell line, p≤0.05, ANOVA with Holm-Sidak post-test

**Figure 14.**
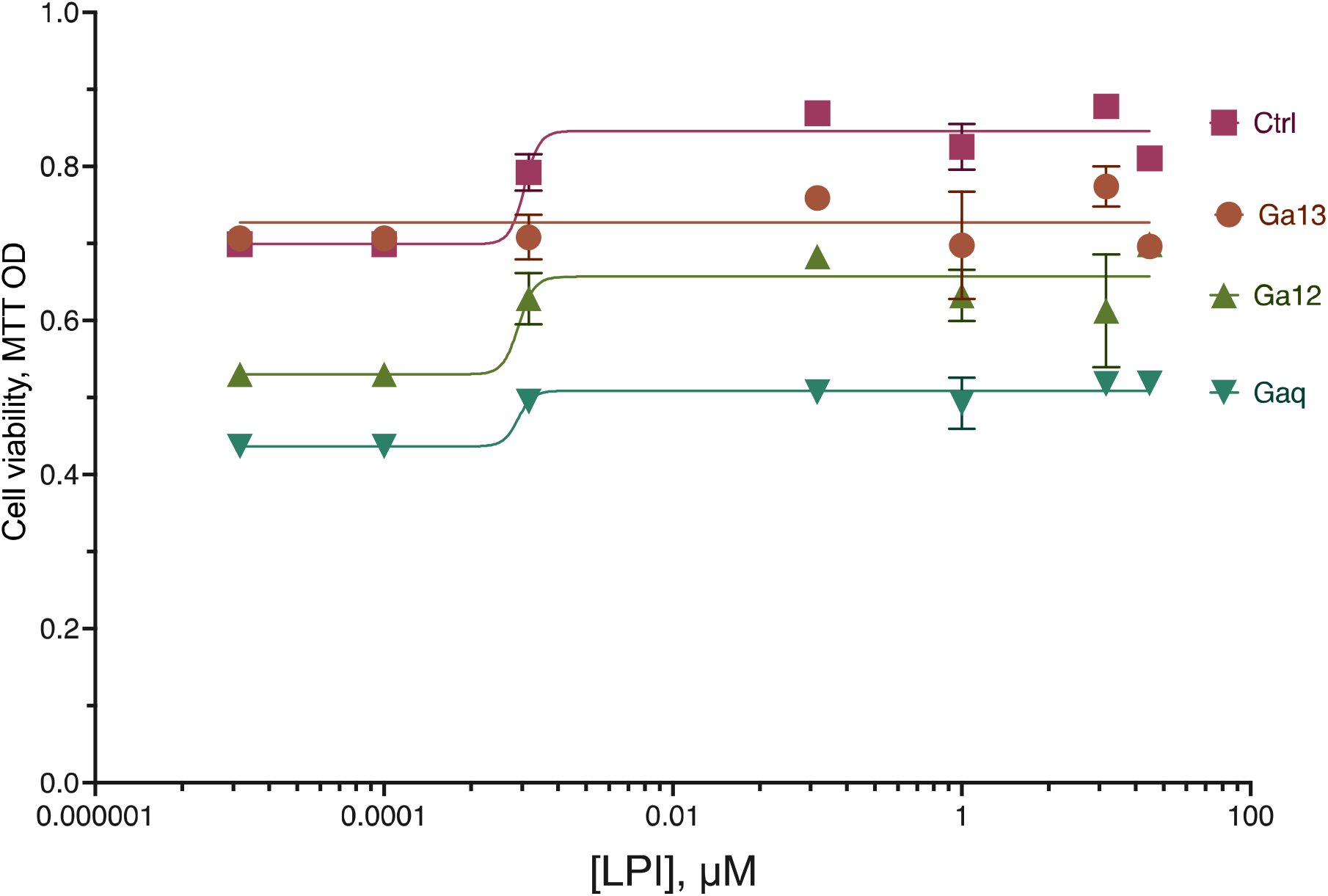
Stimulation of MDA-MB-231 proliferation against the background of knockdown of Gα subunits. Ctrl - scrambled control. Incubation with substance 72 hours. MTT test data, mean±standard deviation

## 3. Discussion

NADA is a CB1 receptor agonist (CB1 Ki = 250 nM) with a 40-fold selectivity over CB2 [9]. At the same time, it acts as a potent TRPV1 activator [10,11].

In this study, the interaction of CB2 and GPR55 receptors was found during the pro-proliferative action of LPI. CB2R and GPR55 form heteromers in cancer cells, that these structures possess unique signaling properties, and that modulation of these heteromers can modify the antitumoral activity of cannabinoids in vivo. via these complexes, agonists and antagonists of one receptor are able to impair the signaling of the partner receptor. At low concentrations, THC (a well reported CB2R agonist) signals through CB2R, producing a conceivable activation of ERK-1/2 and inhibition of FK-induced cAMP increase. At higher concentrations, THC is able to target GPR55, acting as a receptor antagonist and exerting a cross-antagonism over CB2R through the heteromer, which would result in an attenuation of the CB2R-mediated effects on ERK-1/2 activation and cAMP production [12]. Whereas heteromerization leads to a reduction in GPR55-mediated activation of transcription factors (nuclear factor of activated T-cells, NF-κB and cAMP response element), ERK1/2-MAPK activation is potentiated in the presence of CB2 receptors. CB2 receptor-mediated signalling is also affected by co-expression with GPR55 [13].

CB1 and GPR55 receptors are co-expressed and form heteromers in rat and monkey striatum. In the heterologous system, a biochemical fingerprint consisting on cross-antagonism in ERK1/2 phosphorylation was detected. The cross-antagonism was also observed on GPR55-mediated NFAT activation [14]. Within the framework of the model system of this study, the contribution of this heterodimer was not found. It can be assumed that this is due to the direction of action: after the activation of heterodimeric CB1, a weakening of the pro-proliferative action of DHA-DA, which is not typical for this substance, should have followed. It can be assumed that the involvement of this heterodimer can be seen as an increase in the pro-proliferative effect of LPI after CB1 knockdown.

GPR18 is known to form heterodimers with CB2R but not with CB1R. CB2-GPR18 heteroreceptor complexes displayed particular functional properties often consisting of negative cross-talk and cross-antagonism (the response of one of the receptors is blocked by a selective antagonist of the partner receptor) [15]. On the other hand, it is known that pro-apoptotic GPR55 agonists can realize part of their action through the activation of GPR18 [16], which is consistent with the results obtained in this study.

DRG neurons treated with 10 and 50 µMol/L CBD showed calcium influx, but not at lower doses. Neurons treated with capsaicin demonstrated robust calcium influx, which was dose-dependently reduced in the presence of low dose CBD (IC50 = 100 nMol/L). The inhibition or desensitization by CBD was reversed in the presence of forskolin and cyclosporin. Forskolin-stimulated cAMP levels were significantly reduced in CBD treated neurons. CBD at low doses corresponding to plasma concentrations observed physiologically inhibits or desensitizes neuronal TRPV1 signalling by inhibiting the adenylyl cyclase – cAMP pathway, which is essential for maintaining TRPV1 phosphorylation and sensitization. CBD also facilitated calcineurin-mediated TRPV1 inhibition [17]. However, CBD engages different targets and hence CBD’s effects are thought to be due to multiple molecular mechanisms of action [18].

Activation of ERK in primary afferent neurons is mediated, at least in part, by TRPV1 [19].

N-arachidonoyl-dopamine (NADA) and N-oleoyl-dopamine (OLDA), induces concentration-dependent death of PBMC from healthy donors. TRPV1 (5’-iodoresiniferatoxin) and cannabinoid (CB2; AM630) receptor antagonists block the cytotoxic effect of NADA [20]. In other systems, activation of this receptor by acyl dopamines can lead to and increased survival [21].

For the CCKAR receptor, compared to Gα_s_ or Gα_i_ proteins, Gα_q_ coupling increases the binding affinity of CCK-8, consistent with the increased binding activity of isoproterenol against β2AR in the presence of Gα_s_ protein [22]. G-protein binding may allosterically influence ligand binding. G-protein coupling to the β2AR stabilizes a ‘closed’ receptor conformation characterized by restricted access to and egress from the hormone-binding site. In contrast to agonist binding alone, coupling of a G protein in the absence of an agonist stabilizes large structural changes in a GPCR [23].

It can be assumed that within the framework of this study, a similar effect can be realized with a significant drop in the affinity of GPR55 for LPI in the absence of Gα_13_.

## 4. Materials and Methods

### 4.1. Reagents

Isopropanol, MTT, D-glucose, DMSO, RPMI 1640, DMEM, L-glutamine, HCl, Triton X-100, Hank’s salts solution, Versene’s solution, penicillin, streptomycin, amphotericin B, RPMI 1640, DMEM, trypsin, (4,5-dimethylthiazol-2-yl)-2,5-diphenyltetrazolium bromide (MTT), and fetal bovine serum were from PanEco, Moscow, Russia.

Cell lines were purchased from ATCC, Manassas, VA, USA.

Antibodies anti-b-actin and anti-GPR55 were from Abcam, Cambridge, UK. Anti-mouse IgG antibody was from Santa-Cruz Biotechnology, Dallas, TX, US.

666-15, NQDI1, and SP 600125, were from Cayman Europe, Hamburg, Germany. CID 16020046 was from Tocris Bioscience, Bristol, UK. Glycylglycine, acetic acid, MgSO_4,_ EGTA, dithiothreitol, DMSO, Triton X-100, PTIO, *N*-Ac-Cys, cannabidiol, L-NAME, L-NMMA, acrylamide, bis-acrylamide, SDS, nitro blue tetrazolium, Tris-Borate-EDTA, agarose, bicinchoninic acid, D-glucose, bovine serum albumin, anti-rabbit IgG antibody, and 5-Bromo-4-chloro-3-indolyl phosphate-toluoidine were from Sigma-Aldrich, St. Louis, MO, USA. Calcium Green, siRNA, RNAiMax, Advanced DMEM, and DreamTaq master mix were from Thermo Fisher Scientific, Walthon, MA USA. pCREB-Luc CREB luciferase reporter vector was from Signosis, Santa Clara, CA, USA. Total RNA Purification kit was from Jena Biosciences, Jena, Germany. MMLV reverse transcription kit, Red fluorescent protein expression vector pTagRFP-N were from Evrogen, Moscow, Russia. The purity of all used reagents was 95% or more.

### 4.2. Chemical Synthesis

Dopamine amide docosahexaenoic acid [24] was synthesized as described previously.

### 4.3 Cell culture

Cells were cultured in DMEM supplemented with 10% FBS, 4 mM L-glutamine, 100 U/mL penicillin, 100 µg/mL streptomycin, and 2.5 µg/mL amphotericin B in 5% CO2 and 100% humidity at 37°C. Cells were subcultured by successive treatment with Versene’s solution and trypsin solution. Mycoplasma contamination was controlled using the Jena Biosciences kit according to the manufacturer’s instructions.

### 4.4. siRNA knockdown

To test the hypothesis about the involvement of different Gα subunits in the implementation of the cytotoxic and pro-proliferative effects of the GPR55 receptor, the knockdown approach of the Gα_12_, Gα_13_, and Gα_q_ subunits using siRNA was used. For each gene, a commercially available combination of three siRNAs was used, transfection of a commercially available siRNA with a random nucleotide sequence (scrambled) was used as a control.

To optimize the conditions, two variants of transfecting agents were used (Thermo RNAiMax and Thermo Lipofectamine 3000, the latter in two addition variants, 0.75 and 1.5 µl per well of a 24-well plate). For Gα_13_, the use of RNAiMax turned out to be optimal, for the remaining two Gα, Lipofectamine 3000 in variant 20. Under optimal conditions, RNA expression of each of the subunits was not recorded after 72 h of incubation with siRNA.

### 4.5. CRISPR knockdown

Knockout lines were generated using the CRISPR-Cas9 system described by Ran et al.[25].

At the first stage, plasmid constructs based on the PX459 vector (pSpCas9(BB)-2A-Puro) containing sequences of 20 nucleotides complementary to the modified genomic DNA region were obtained. A requirement for the selection of Cas9 target sites is the presence of a PAM sequence located immediately 3’ of the 20 bp target sequence. Each Cas9 orthologue has a unique PAM sequence; for example, SpCas9 requires 5’-NGG. For the convenience of primary screening, the target sequences were chosen so that, in addition to the 5’-NGG sequence, at the 3’-end they contained cleavage sites with restriction enzymes; in the case of successful genomic modification, one or more nucleotides are excised and the restriction site disappears. To obtain GPR55 knockout cells into the PX459 vector at the BbsI restriction sites, oligonucleotides were cloned that were complementary to the modified Gpr55fw gene region, 5’-CACCGTCCCTATCTACAGTTTCCAT-3’ and Gpr55rev, 5’-AAACATGGAAACTGTAGATAGGGAC-3’. The genomic DNA contained the NcoI restriction site at the Cas9 cleavage site. Sequencing of the resulting plasmid confirmed the presence of the target insert.

Next, the MDA-MB-231 cell line was transfected with the obtained plasmid DNA using Lipofectamine 3000 (Invitrogen) according to the manufacturer’s protocol. Two days after transfection, cells were selected on puromycin, an antibiotic was added to the cells at a concentration of 1 to 3 µg/ml, and the cells were incubated for three days. Non-transfected MDA-MB-231 cells were used as controls. After selection on puromycin, genomic DNA was isolated from the cells, and the gene region containing the target modifiable sequence was amplified with primers hGpr55_check_fw – 5’-GGTGGAGTGCCTTTTACTTCGTCAGC-3’ and hGpr55_check_rev – 5’-TCTGCTGCACCCAGTCCTGGGTGTG-3’.

To obtain CNR1 knockout cells, oligonucleotides were cloned into the PX459 vector at the BbsI restriction sites, complementary to the regions of the modifiable CNR1rev genes, 5’-AAACGCAGCGGAGGCTGCGGGAGTC-3’ and CNR1fw, 5’-CACCGACTCCCCGCAGCCTCCGCTGC-3’. For CNR2, primers CNR2rev – 5’-AAACAGGGTCACCAGTGCCCTTCCC-3’ and CNR2fw – 5’-CACCGGGAAGGGCACTGGTGACCCT-3’ were used.

For knockout detection, primers testCNR1_fw 5’-GAGAACTTCATGGACATAGAGTG-3’ and test CNR1_rev 5’-GGAGGCCGTGACCCCACCCAGTT-3’ for gene CNR1, CNR2 test_fw 5’-TGTCTTCCTGCTGAAGATTGGCAG-3’ and CNR2 test_rev 5’-CGGAAAAGAGGAAGGCGATGAACAG-3’

Obtaining a cell line knockout for the CNR1 gene. Oligonucleotides were cloned into the PX459 vector at the BbsI restriction sites, complementary to the region of the modifiable CNR1_fw gene, 5’-CACCGCCTGAACCCCAGCCAGCAGC-3’ and CNR1_rev, 5’-AAACGCTGCTGGCTGGGGTTCAGGC-3’. CNR1 genomic DNA contained the PvuII restriction site at the Cas9 cleavage site.

After selection on puromycin, genomic DNA was isolated from the cells, and the gene region containing the target modifiable sequence was amplified. For this, primers CNR1_test_fw 5’-AGATGACTGCGGGAGACAACC-3’ and CNR1_test_rev 5’-CACGTGGAAGTCAATGAAGCTG-3’ were used.

To obtain the TRPV1 gene knockout MDA-MB-231 cell line, oligonucleotides complementary to the region of the modifiable TRPV1_fw gene, 5’-CACCGCGTCCAGGCTGCGGCCCATG-3’ and TRPV1_rev, 5’-AAACCATGGGCCGCAGCCTGGACGC-3’, were cloned into the PX459 vector at the BbsI restriction sites. The TRPV1 genomic DNA contained the NcoI restriction site at the Cas9 cleavage site. Next, the MDA-MB-231 cell line was transfected with the obtained plasmid DNA using Lipofectamine 3000 (Invitrogen) according to the manufacturer’s protocol. Two days after transfection, cells were selected on puromycin, the antibiotic was added to a final concentration of 1 to 3 µg/ml, and the cells were incubated for three days. After selection on puromycin, genomic DNA was isolated from the cells, and the gene region containing the target modifiable sequence was amplified. For this, primers TRPV1_test_fw – 5’-GAGATCCAGCTGGCCCCTG-3’ and TRPV1_test_rev – 5’-GAGCCCTTCCCCAGCACCT-3’ were used.

### 4.6. RT-qPCR

Real-time PCR with gene-specific primers was used to control the effectiveness of knockdown:

Ga12 straight 5’-GGGCGAGTGAAACTGAAA-3’, reverse 5’-CACAACACGGTCCTCAATTAAAC-3’

Ga13 straight 5’-CACTGCTTAAGAGACGTCCAA-3’, reverse 5’-CAGTGGTGAAGTGGTGGTATAA-3’

Gaq direct 5’-GCCACAGACACCGAGAATATC-3’, reverse 5’-GGTGTCTAGGAGGCACAATTAG-3’.

For each pair of primers, it was mandatory to check nonspecific amplification using RNA without adding reverse transcriptase at the stage of cDNA production.

### 4.7. RNA Isolation and RT-PCR

Total RNA was isolated using the Total RNA Purification Kit (Jena Biosciences) according to the manufacturer’s protocol. Residual genomic DNA was removed using DNase I (Thermo Fisher Scientific, Walthon, MA USA) according to the manufacturer’s protocol; 1 U of the enzyme was used per RNA sample. cDNA was synthesized using the MMLV reverse transcription kit (Evrogen, Moscow, Russia) with an oligo-dT primer. PCR was performed using the DreamTaq master mix (Thermo Fisher Scientific, Walthon, MA USA); the program was as follows. Initial denaturation at 95 °C for 3 min, cycle: denaturation at 95 °C for 30 s, annealing at 57 °C for 30 s, DNA synthesis at 72 °C for 30 s for 35 cycles, final DNA synthesis at 72 °C for 5 min. The primers were generated using the ITDNA PrimerQuest tool (https://eu.idtdna.com/PrimerQuest; accessed on 01.01.2020) and validated using the NCBI Primer-BLAST service [26].

### 4.8. Cytotoxicity evaluation

Cells were seeded in 96-well plates in the amount of 15 thousand per well in a volume of 100 µl of medium and cultured for a day. After that, the substance was added at the required concentration in the range from 0.01 to 150 µM in the form of a DMSO solution in an additional 100 µl of the medium; the final DMSO concentration was 0.5% or less. In the event that receptor blockers were used, they were added to 50 µl of the medium one hour before the addition of the substance and then the second portion (to ensure the constancy of concentration) with the substance; the volume of the medium in which the substance was added was 50 µl in this case. The cells were incubated with the substance for 24 hours, after which the viability was determined using the MTT test. To do this, the culture medium in the wells was replaced with an MTT solution in Earl’s solution with the addition of 1 g/L D-glucose and incubated for 1.5 hours at 37C under cell culture conditions. After that, an equal volume of 0.04 M HCl in isopropanol was added to the wells and intensively stirred at 37C on a shaker for 30 minutes. At the end, the optical density of the solution was determined at a wavelength of 570 nm with subtraction of the background at a wavelength of 660 nm. Each experiment was repeated at least five times.

### 4.9. Western Blot

To evaluate the expression of particular proteins in the cells, the cells were seeded at the density 200,000 per well of 24-well plate the day before experiment. After the appropriate treatment, the cells were washed once with PBS, lysed using the lysis solution (150 mM NaCl, 1% Triton X-100, 0.1% SDS, 50 mM Tris-HCl pH 8.0, 1% protease inhibitor cocktail) for 30 min at +4 °C, and centrifuged for 5 min at 10,000× g. Total protein concentration in the supernatants was determined using the BCA assay. Proteins were separated using denaturing SDS-PAGE in 10% gel with the PageRuler protein ladder, transferred to a nitrocellulose membrane using the Invitrogen Power Blotter with the Invitrogen Power Blotter 1-step transfer buffer and Invitrogen precut membranes and filters, and stained with antibodies using the Invitrogen iBind system according to the manufacturer’s protocol. The following antibodies were used: rabbit anti-GPR55 (Abcam ab203663), mouse anti-beta-actin (Abcam ab8226); secondary antibodies (coupled to alkaline phosphatase) anti-rabbit IgG (Sigma-Aldrich A9919), anti-mouse IgG (Santa-Cruz Biotech scbt-2008). After the staining, the membrane was washed in H_2_O for 10 min and incubated with the staining solution (20 µL BCIP solution + 30 µL NBT solution per 10 mL of substrate buffer) for 1 h at room temperature. Substrate buffer for alkaline phosphatase: 100 mM Tris-HCl, pH 9.5, 100 mM NaCl, and 5 mM MgCl_2_. BCIP solution: 20 mg/mL 5-Bromo-4-chloro-3-indolyl phosphate-toluoidine (BCIP) in 100% di-methyl formamide. NBT staining solution: 50 mg/mL nitro blue tetrazolium (NBT) in 70% di-methyl formamide.

### 4.10. BCA Protein Assay

Protein concentration was determined using the BCA assay [80]. The following base reagents were used: Reagent A (bicinchoninic acid 1%, Na_2_CO_3_*H_2_O 2%, sodium tartrate 0.16%, NaOH 0.4%, NaHCO_3_ 0.95%, pH 11.25), Reagent B (4% CuSO_4_*5H_2_O), S-WR (50 volumes of Reagent A + 1 volume of Reagent B). 5 µL of cell lysate was mixed with 40 µL of S-WR and incubated for 15 min at 60 °C, after which the optical density was measured at λ = 562 nm using the Hidex Sense Beta Plus microplate reader (Hidex, Fineland). Each sample was assayed in triplicate. Cell lysis buffer was used as a background control. Bovine serum albumin solution in the cell lysis buffer was used as a positive control and to build a calibration curve.

### 4.11. Statistical analysis

Statistical evaluation was performed using the GraphPad Prism 9.3 software. ANOVA with the Holm-Sidak post-test was used to compare the obtained values; p≤0.05 were considered significant.

## 5. Conclusions

Thus, the GPR18 and TRPV1 receptors are additionally involved in the implementation of the cytotoxic effect of DHA-DA, while the CB1 receptor is not involved. The data obtained are consistent with the literature data on the possibility of GPR18 involvement in the cytotoxic effect of GPR55 ligands, as well as on the ability of vaniloids to induce cell death through overactivation of the TRPV1 cation channel. When proliferation is stimulated with the participation of the GPR55 receptor, a signal is transmitted from the CB2 receptor to the GPR55 receptor due to the formation of a heterodimer. In the implementation of the cytotoxic action of DHA-DA, the predominant participation of one of the Gα subunits was not found, but the Gα_13_ subunit plays a decisive role in the implementation of the proproliferative action.

## Supplementary Materials

Not applicable.

## Author Contributions

Conceptualization, M.G.A. and V.V.B.; methodology, M.G.A., N.M.G., and I.E.D..; investigation, A.M.G., P.V.D., N.M.G., G.D.S., K.O.D., O.V.S., and G.N.Z.; writing—original draft preparation, M.G.A. and O.V.S.; writing—review and editing, V.V.B.; visualization, M.G.A. and O.V.S.; supervision, V.V.B. and I.E.D.; project administration, V.V.B., I.E.D. and M.G.A.; funding acquisition, V.V.B. All authors have read and agreed to the published version of the manuscript.

## Institutional Review Board Statement

Not applicable.

## Informed Consent Statement

Not applicable.

## Data Availability Statement

The data presented in this study are available on request from the corresponding author. The data are not publicly available due to legal issues.

## Acknowledgments

Not applicable.

## Abbreviations

DHA-DA: N-docosahexaenoyl dopamine
LPI: lysophosphatidylinositol
ROS: reactive oxygen species
FAAH: fatty acid amide hydrolase
CBD: cannabidiol
NADA: N-acyldopamines
BCIP: 5-Bromo-4-chloro-3-indolyl phosphate
NBT: nitro blue tetrazolium
BCA: bicinchoninic acid

